# Variant Analysis Pipeline for Accurate Detection of Genomic Variants from Transcriptome Sequencing Data: SNP Calling from RNA-seq Data in Non-Human Models

**DOI:** 10.1101/625020

**Authors:** Modupeore O. Adetunji, Carl J. Schmidt, Susan J. Lamont, Behnam Abasht

## Abstract

The wealth of information deliverable from transcriptome sequencing (RNA-seq) is significant, however current applications for variant detection still remain a challenge due to the complexity of the transcriptome. Given the ability of RNA-seq to reveal active regions of the genome, detection of RNA-seq SNPs can prove valuable in understanding the phenotypic diversity between populations. Thus, we present a novel computational workflow named VAP (Variant Analysis Pipeline) that takes advantage of multiple RNA-seq splice aware aligners to call SNPs in non-human models using RNA-seq data only. We applied VAP to RNA-seq from a highly inbred chicken line and achieved >97% precision and >99% sensitivity when compared with the matching whole genome sequencing (WGS) data. Over 65% of WGS coding variants were identified from RNA-seq. Further, our results discovered SNPs resulting from post translational modifications, such as RNA editing, which may reveal potentially functional variation that would have otherwise been missed in genomic data. Even with the limitation in detecting variants in expressed regions only, our method proves to be a reliable alternative for SNP identification using RNA-seq data.

## Introduction

Detection of single nucleotide polymorphisms (SNPs) is an important step in understanding the relationship between genotype and phenotype. The insights achieved with next generation sequencing (NGS) technologies provide an unbiased view of the entire genome, exome or transcriptome at a reasonable cost (1). Most methods for variant identification utilize whole-genome or whole-exome sequencing data, while variant identification using RNA-seq remains a challenge because of the complexity in the transcriptome and the high false positive rates (2). However, having access to RNA sequences at a single nucleotide resolution provides the opportunity to investigate gene or transcript differences across species at a nucleotide level.

RNA-seq is applicable to numerous research studies, such as the quantification of gene expression levels, detection of alternative splicing, allele-specific expression, gene fusions or RNA editing (3). Workflows have been developed to address identifying SNPs from RNA-seq reads in human, including SNPiR and eSNV-detect. SNPiR (4) employs BWA aligner and variant calling using GATK UnifiedGenotyper, eSNV-detect (5) relies on combination of two aligners (BWA and TopHat2) followed by variant calling with SAMtools and Opposum + Platypus (6). Opposum reconstructs RNA alignment files to make them suitable for haplotype-based variant calling with Platypus (7). These workflows require adequate sampling of RNA-seq reads and accurate mapping of the RNA-seq reads to the reference genome to avoid false positive SNP calls. In addition to the limitation of these workflows being specifically designed for human samples, they either rely on outdated variant calling procedures, or preprocessing RNA-seq data to make it suitable for variant calling, thus making it difficult to sufficiently compare their performance.

Due to the aforementioned limitations, we designed a workflow, called VAP (Variant Analysis Pipeline), to reliably identify SNPs in RNA-seq in non-human models. VAP takes into consideration current state-of-the-art RNA-seq mapping, variant calling algorithms and the GATK best practices recommended by the Broad Institute (8), Our workflow consists of (i) multiple splice-aware reference-mapping algorithms that make use of the transcripts annotation data, (ii) variant calling following the Genome Analysis Toolkit (GATK) best practices, and (iii) stringent filtering procedures. We propose that calculating specificity will estimate the likelihood of detecting a true variant in RNA-seq and sensitivity will determine how likely RNA-seq is able to detect an expressed SNP if it is present in a transcribed gene (9). Overall the results indicate that RNA-seq can be an accurate method of SNP detection using our VAP workflow.

## Materials and methods

### VAP Workflow

Fig 1 shows the flowchart of the VAP workflow. Read quality was assessed using FastQC and preprocessed using Trimmomatic (10) and/or AfterQC (11) when required. Pre-processed RNA-seq reads were mapped to the reference genome and known transcripts employing three splice-aware assembly tools; TopHat2 (12), HiSAT2 (13) and STAR (14). All three programs are open-source and are highly recommended for reliable reference mapping of RNA-seq data (15). SAMtools was used to convert the alignment results to BAM format (16). The mapped reads undergo sorting, adding read groups, and marking of duplicates using Picard tools package (https://broadinstitute.github.io/picard/). The SNP calling step uses the GATK toolkit for splitting “N” cigar reads (i.e. splice junction reads), base quality score recalibration and variant detection using the GATK HaplotypeCaller (17). Lastly, the filtering steps entail assigning priority to SNPs found in all three mapping plus SNP calling steps, to minimize false positive variant calls. The priority SNPs were filtered using the GATK Variant Filtration tool and custom Perl scripts. SNPs were filtered using the set of read characteristics summarized in Table 1; low quality calls (QD < 5), or variants with strong strand bias (FS > 60), or low read depth (DP < 10) and SNP clusters (3 SNPs in 35bp window) were excluded from further analysis. Custom filtering was described as follows: nucleotide positions with less than 5 reads supporting alternative allele and nucleotide positions with heterozygosity scores < 0.10 are eliminated to prevent ambiguous SNP calls. Alternative-allele ratio *(Het)* is calculated by *Heti* = *aa_i_ / t_i_*; where *i* is the nucleotide base pair, *aai* is the alternate read depth at the location *i*, and *t_i_* is the total number of reads at location *i*. After filtering, the variants were annotated using the ANNOVAR (18) and VEP (19) software.

**Fig 1.**
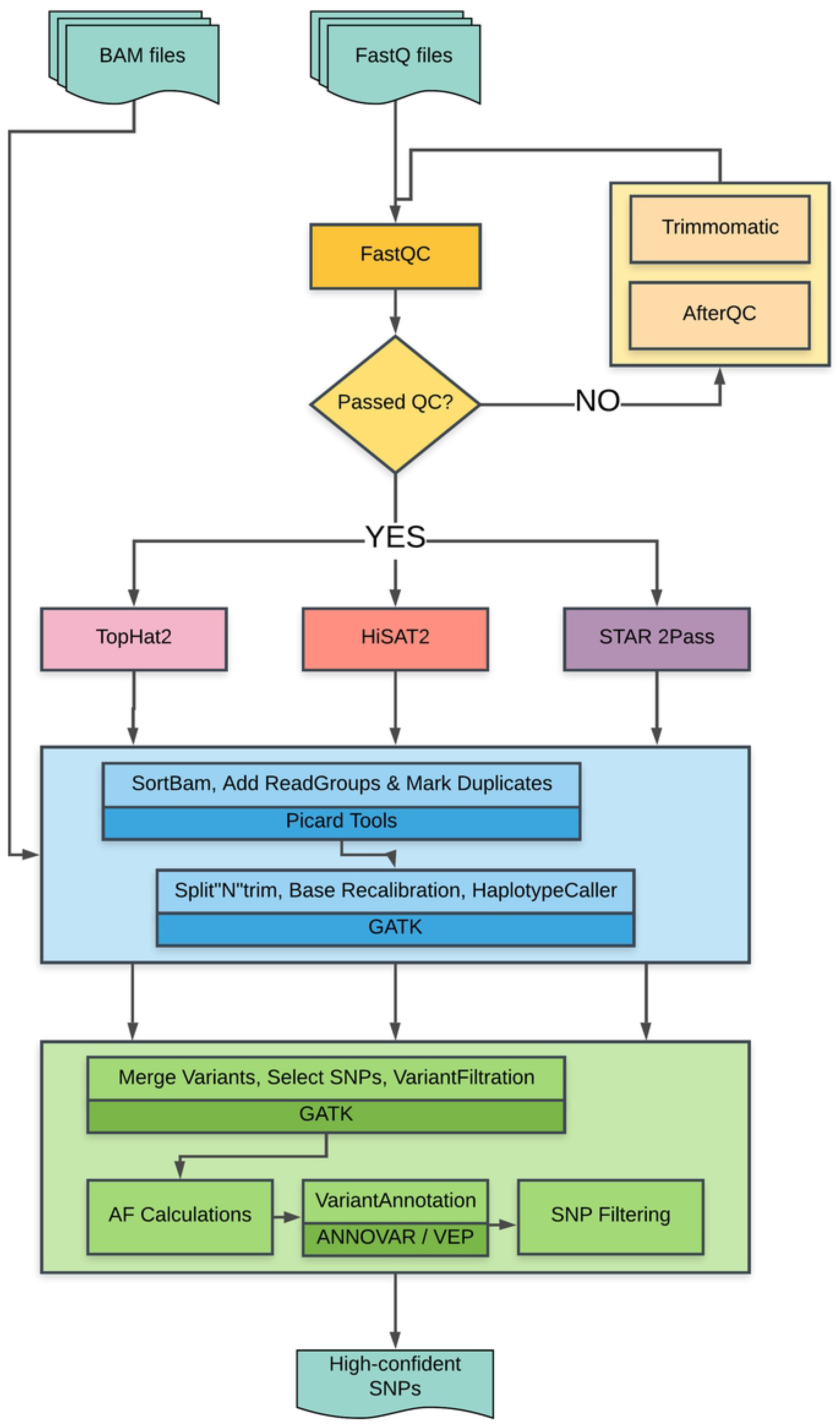
Flow chart of the VAP workflow. FastQ files are QC using FastQC, mapped using three aligners. BAM files are pre-processed by Picard and GATK, then merged, annotated and filtered to achieve high-confident SNPs.

**Table 1.** Criteria used in the VAP filtering workflow.

| Criteria | Threshold |
| --- | --- |
| GATK - VariantFiltration tool |  |
| ReadRankPosSum (RRPS) | $RRPS < -8$ |
| Quality by depth (QD) | $QD < 5$ |
| Read depth (DP) | $DP < 10$ |
| Fisher's exact test p-value (FS) | $FS > 60$ |
| Mapping Quality (MQ) | $MQ < 40$ |
| SnpCluster | 3 SNPs in 35bp |
| Mann-Whitney Rank-Sum (MQRankSum) | MQRankSum < -12.5 |
| Alternative allele supporting read depth | ALTreads < 5 |
| Alternative allele ratio (*Het*) | $aa / t \leq 0.10$ |

### DNA and RNA Sequencing data

We obtained RNA-seq and whole genome sequencing (WGS) data for highly inbred Fayoumi chickens from previously published works. For RNA-seq, samples were collected from the brain and liver generating of 2 chicken embryos at day 12, generating 117 million 75bp pair-end reads (Zhuo et al., 2017; the NCBI Sequence Read Archive Accession number SRP102082) (20). For WGS, pooled DNA samples were constructed from individual DNA isolates from blood from 16 birds, contributing to 241 million 100bp pair-end reads (Fleming et al., 2016; the NCBI Sequence Read Archive Accession number SRP192622) (21). Both samples were sequenced on the Illumina HiSeq platform. The transcriptome and whole genome of these samples have been deeply sequenced to provide sufficient coverage for accurate identification of variants from RNA and DNA of the same line. Having matched RNA and DNA samples allows for suitable verification of RNA SNP calls, making our datasets good candidates for evaluating the accuracy of our VAP methodology.

### 600K Genotyping data

Samples were genotypes individually and included 96 samples from two purebred (23 samples) and one crossbred (73 samples) commercial broiler populations. The samples were genotyped with the ThermoFisher Axiom Chicken Genotyping Array (22). The raw genotyping data (cel files) was analyzed with the *Gallus gallus* 5.0 genome (from Axiom server) using the Axiom Analysis Suite Software (version 3.0.1) following the software’s Best Practices Workflow using recommended settings for agricultural animals. The final results were exported, including a raw VCF of all the genotype calls and a *txt* file of all variants with >= 97% call rate. The *txt* file was utilized to filter low quality variants from the raw VCF.

### RNA-seq Mapping, Variant Calling and Filtering

RNA-seq samples were mapped with the three RNA-seq mapping tools; TopHat2 (v 2.1.1), HiSAT2 (v 2.1.0) and STAR (v 2.5.2b) 2-pass method using default parameters to the NCBI *Gallus gallus* Build 5.0 reference genome and the mapping files were converted to BAM using SAMtools (v 1.4.1). The BAM files were processed, and variants were called using Picard tools (v 2.13.2) and GATK (v 3.8-0-ge9d806836) through the VAP pipeline. We used ANNOVAR (v 2017Jul16) and VEP (v 91) to annotate variants on the basis of gene model from RefSeq, Ensembl and the UCSC Genome Browser. We retained SNPs found with all three mapping tools and those that fulfilled the filtering criteria in Table 1. SNPs found in WGS data or present in dbSNP (Build 150) are identified as “verified” variants, while those not found are tagged as “novel”. The precision of the VAP workflow was determined as the number of all known RNA-seq variants divided by the total number of known and novel RNA-seq variants, i.e. Precision = verified_SNPs_ / (verified_SNPs_ + novelSNPs).

### WGS Mapping, Variant Calling and Filtering

We mapped the WGS data with BWA-mem (v 0.7.16a-r1181) (23) using default parameters to the NCBI *Gallus gallus* Build 5.0 reference genome. Variant calling was performed using Picard and GATK HaplotypeCaller, following the recommendations proposed by Van der Auwera et al (24) and Yiyuan Yan et al (25). Similar filtering parameters for RNA-seq as previously described were applied using the GATK Variant Filtration tool and custom scripts (Table 1). To allow a fair comparison between RNA-seq and WGS variants, we estimated specificity with the fraction of coding exonic variants identified from WGS.

### Sensitivity and Specificity of Verified RNA-seq SNPs

To determine the accuracy of detecting a true variant from RNA-seq using our VAP workflow, we calculated the specificity and sensitivity of the verified RNA-seq SNPs. Because we are using transcriptome data, we should only be theoretically able to detect SNPs at sites expressed in our data. Sensitivity analysis will evaluate the accuracy of our pipeline to correctly detect known SNPs using RNA-seq, and specificity analysis will assess how likely a SNP is detected by RNA-seq compared to WGS. To do this, we further characterized our verified RNA-seq SNPs as “true-verified” and “non-verified” SNPs. A true-verified SNP (TS) is a SNP with the same corresponding dbSNP and/or WGS data, and a non-verified SNP (NS) is where the genotype does not match the dbSNP/WGS data. Also, SNPs not detected in RNA-seq but found in WGS and validated using dbSNP are called “DNA-verified” SNPs (DS). Sensitivity is calculated as the number of TS divided by the number of TS plus the number of PS (i.e. Sensitivity = TS / (TS + NS)). While specificity is estimated as the number of TS divided by the number of TS plus the number of DS (i.e. Specificity = TS / (TS + DS)) (4,9).

### Gene Expression Analysis

Variants in expressed regions were identified by gene quantification analysis using StringTie v1.3.3 (26) on the TopHat2, HISAT2 and STAR BAM files. The average FPKM (fragments per kilobase of transcript per million fragments mapped) was calculated for specificity analysis.

## RESULTS

### The Multi-Aligner Concept

VAP uses a multi-aligner concept to call SNPs confidently. The application of multiple aligners reduces false discovery rates significantly, as shown in the eSNV-detect pipeline (5,27). However, we do not assign a confidence hierarchy on candidate SNP calls, rather SNP detected from all three aligners are weighted equally, thus all consensus SNPs are obtained and filtered based on the filtering criteria listed above. High percentages of similar SNPs were observed between all three tools, which shows that using a splice-aware read mapper is appropriate for reference mapping using RNA-seq, unlike with BWA. Table 2 provides the summary of mapping and variant calling statistics from the multiple aligners.

**Table 2.** Summary from the multiple aligners; read mapping statistics and variant calls.

| Tools | % reads mapped | % reference covered | Variants | SNPs | % similar SNPs |
| --- | --- | --- | --- | --- | --- |
| TopHat | 87.7 | 23.07 | 578655 | 535505 | 96.12 |
| HiSAT | 90.53 | 23.44 | 636948 | 583547 | 88.21 |
| STAR | 87.81 | 23.7 | 798696 | 708391 | 72.66 |

### SNPs detected in RNA-seq data

Our method identified 514,729 SNPs from all 3 aligners before filtering, which assures reduction of false positives calls (Fig 2). After filtering, 282,798 (54.9%) high confidence SNPs remain, of which 97.2% (274,777 SNPs) were supported by evidence from WGS or dbSNP v.150 (Fig 3). The verified sites exhibited a transition-to-transversion (ts/tv) ratio of 2.84 and estimated ts/tv ratio of ~5 for exonic regions and thus a good indicator of genomic conservation in transcribed regions. For the remaining (novel) 8,021 SNPs, we observed slightly lower ts/tv ratio (2.81) than for the verified sites. The variant sites showed a clear enrichment of transitions, inclusive of A>G and T>C mutations (73.9%), indicative of mRNA editing and the dominant A-to-I RNA editing (28) (Fig 4).

**Fig 2.**
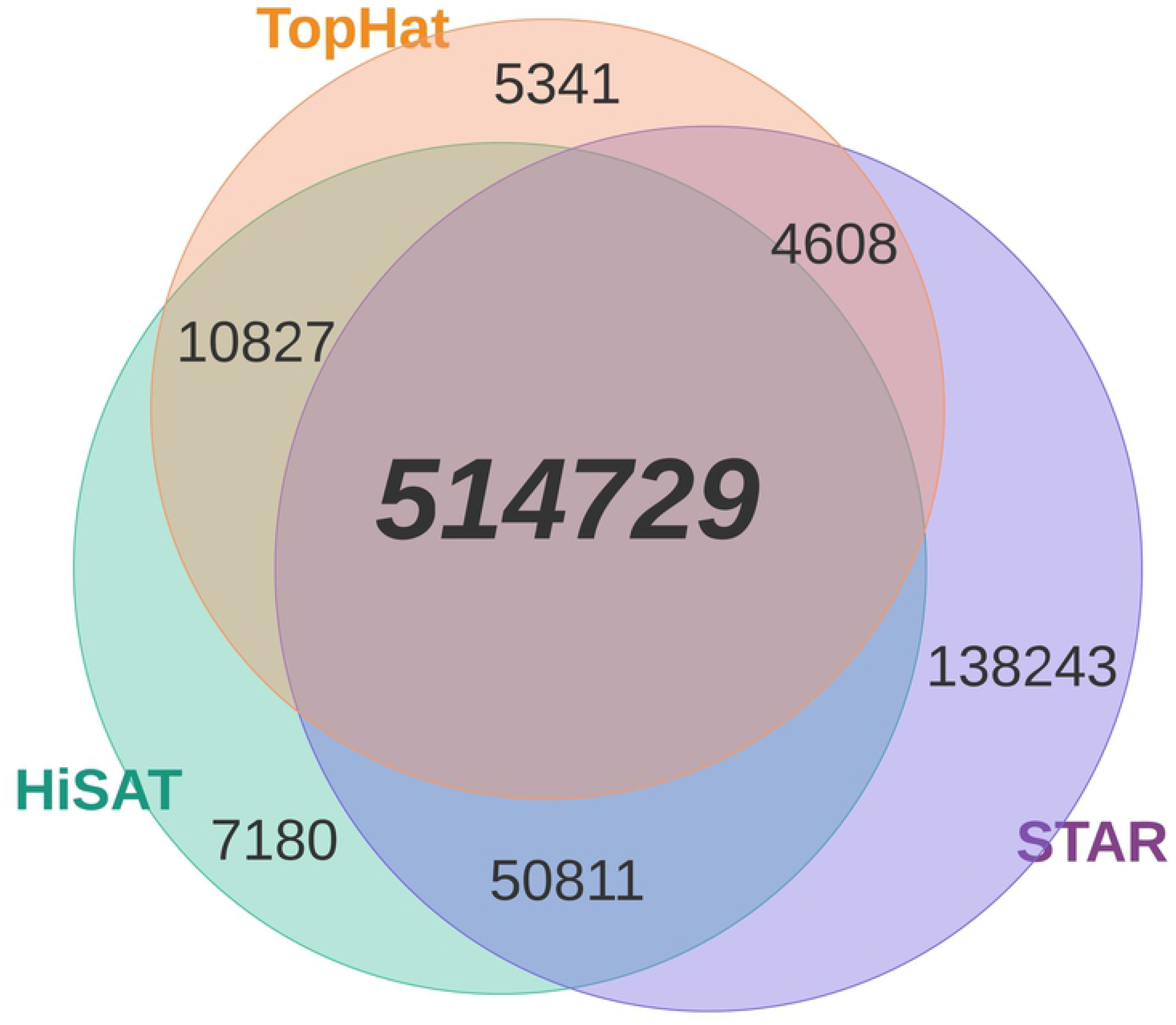
Comparison of RNA-seq SNPs Identified in the different mapping tools.

**Fig 3.**
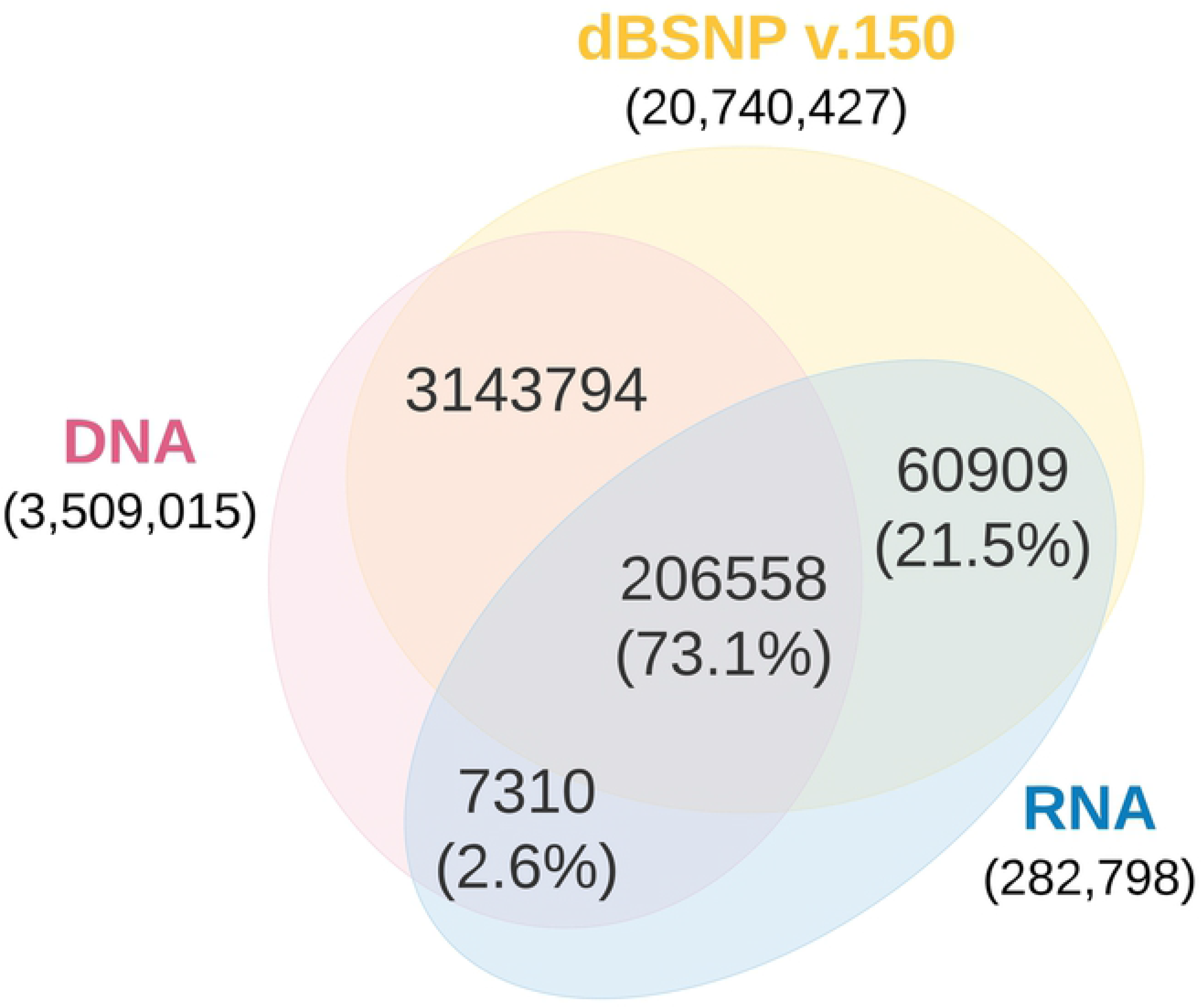
Comparison of RNA-seq SNPs found in either dbSNP or WGS.

**Fig 4.**
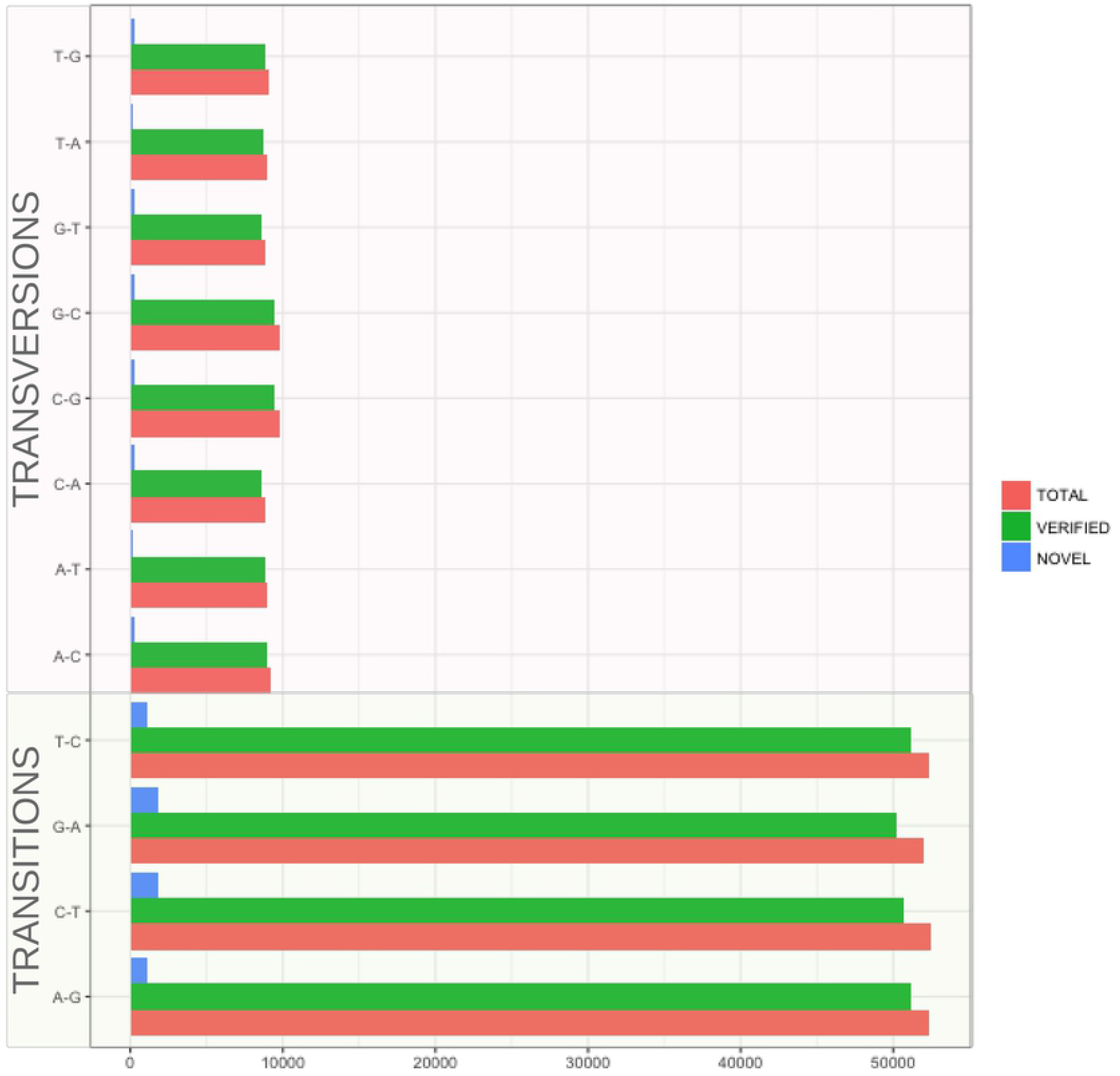
The mutational profile of RNA-seq variants.

### SNPs Allele Frequencies

The 282,798 SNPs called, were grouped based on their variant allele frequencies (VAF). VAFs were calculated by dividing the number of reads supporting the variant allele by the total number of reads obtained. SNPs were grouped as homozygous to the alternative allele with *VAF* ≥ 0.99, and heterozygous with *VAF* < 0.99. We found 264,790 (93.6%) and 18,008 (6.4%) SNPs were classified as homozygous alternate and heterozygous, respectively. Not surprisingly, most of the predicted SNPs were homozygous to the non-reference allele, suggesting the large genetic difference of the Fayoumi breed compared to the reference genome *Gallus gallus* (Red Jungle Fowl) is influenced by polymorphisms (29,30). This will aid in identifying the genomic regions/loci enriched by selection.

### Precision and Sensitivity of RNA-seq SNPs

A high proportion of SNPs detected in RNA-seq data are true variants. The sensitivity of SNP calls are similar for both heterozygous and homozygous sites (Fig 5). With the high number of calls verified via dbSNP, the precision is much higher for homozygous variants compared to heterozygous variants, indicating that a high proportion of expected variants can be detected using RNA-seq with adequate coverage. The decreased precision in heterozygous SNPs may suggest expression of the non-reference allele, and this provides the opportunity to study the effects of genetic variation on the different transcriptional events, such as RNA editing, alternate splicing and allelic specific expression, which cannot be explained using DNA sequencing data (31).

**Fig 5.**
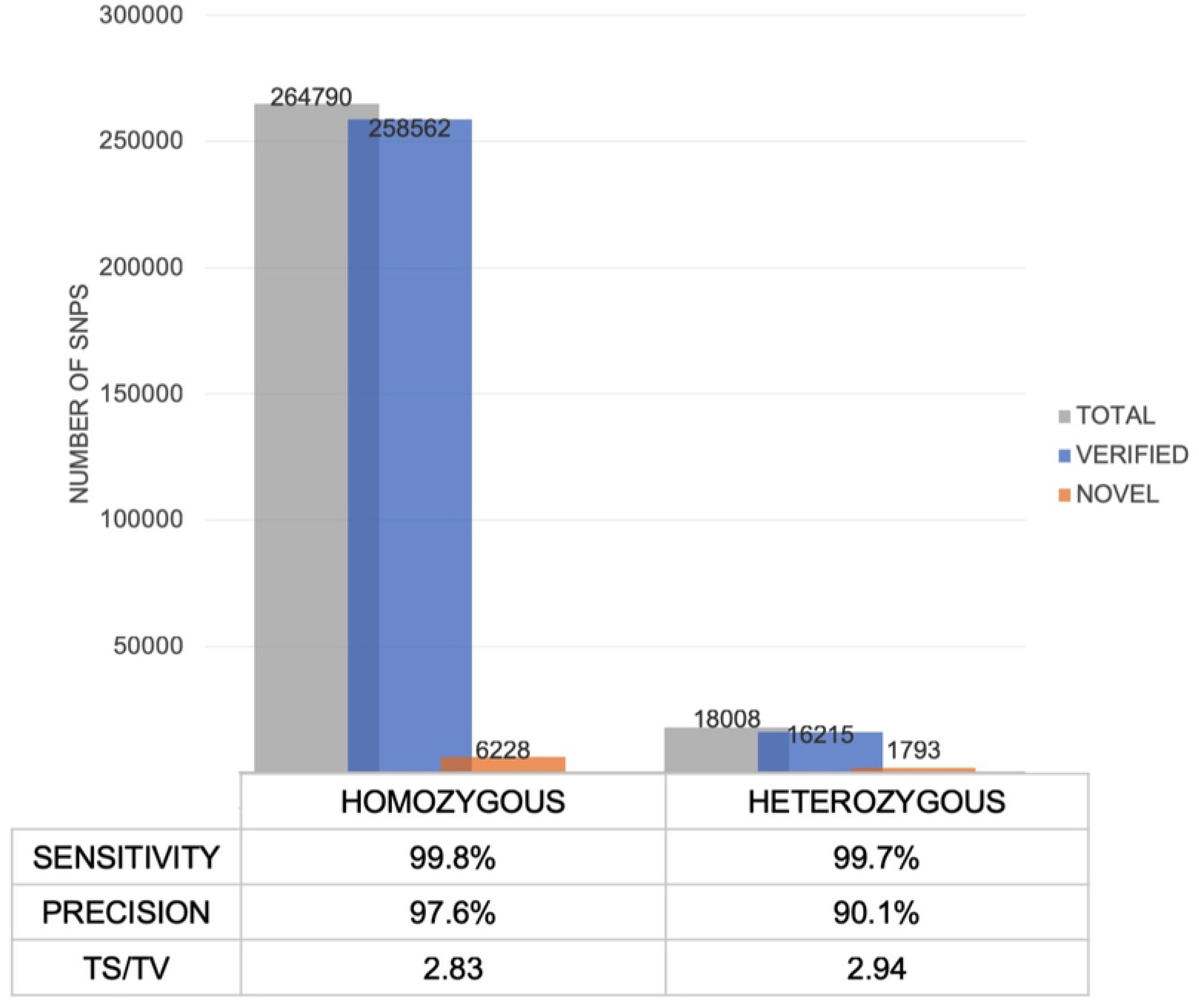
Comparison of SNPs identified as homozygous and heterozygous in RNA-seq.

### Functional classification of RNA-seq and WGS variants

Thirteen percent of the RNA-seq SNPs were predicted to be within protein-coding regions while >1% of the WGS SNPs were in coding regions when annotated against both the NCBI and ENSEMBL gene database for chicken; the remaining SNPs were found in non-coding or regulatory regions (Table 3). Due to difficulty in annotating and determining the impact of polymorphisms on non-coding or regulatory regions, only polymorphisms found on coding regions were further evaluated.

**Table 3.** SNPs belonging to different annotation categories.

| | Annotation categories | Number (%) | Mean VAF (± SD) | No. homozygous (VAF $\geq 0.99$)<sup>a</sup> |
| --- | --- | --- | --- | --- |
| RNA | Intergenic | 162240 (57) | 0.99 (0.06) | 152732 (94%) |
|  | Up/downstream | 11793 (4) | 0.99 (0.07) | 10817 (92%) |
|  | Intronic | 58028 (20) | 0.99 (0.05) | 55744 (96%) |
|  | Exonic | 36702 (13) | 0.99 (0.08) | 33051 (90%) |
|  | Non-synonymous | 8599 (3) | 0.98 (0.11) | 7664 (89%) |
|  | Synonymous | 28094 (10) | 0.99 (0.07) | 25353 (90%) |
|  | Stop-gain/loss | 39 (<1) | 0.96 (0.16) | 34 (87%) |
|  | Splicing | 8 (<1) | 1 (0) | 8 (100%) |
|  | UTR3/UTR5 | 13421 (5) | 0.98 (0.09) | 11895 (88%) |
|  | ncRNA | 106 (<1) | 0.97 (0.13) | 100 (94%) |
| **WGS** | Intergenic | 2865498 (82) | 0.99 (0.07) | 2659382 (92%) |
|  | Up/downstream | 30741 (<1) | 0.99 (0.08) | 28558 (93%) |
|  | Intronic | 565323 (16) | 0.99 (0.07) | 522577 (92%) |
|  | Exonic | 34294 (1) | 0.98 (0.09) | 31875 (92%) |
|  | Non-synonymous | 8946 (<1) | 0.97 (0.11) | 8283 (86%) |
|  | Synonymous | 25274 (<1) | 0.99 (0.08) | 23526 (93%) |
|  | Stop-gain/loss | 74 (<1) | 0.98 (0.11) | 66 (69%) |
|  | Splicing | 17 (<1) | 0.97 (0.13) | 17 (100%) |
|  | UTR3/UTR5 | 12476 (<1) | 0.99 (0.07) | 11515 (92%) |
|  | ncRNA | 302 (<1) | 0.99 (0.07) | 277 (91%) |
| **Overlap RNA & WGS<sup>b</sup>** | Intergenic | 125218 (58) | 1 (0.04) | 112462 (89%) |
|  | Up/downstream | 9787 (4) | 0.99 (0.04) | 6908 (87%) |
|  | Intronic | 47894 (22) | 1 (0.04) | 43636 (91%) |
|  | Exonic | 22551 (10) | 0.99 (0.05) | 19533 (87%) |
|  | Non-synonymous | 5165 (2) | 0.99 (0.06) | 4486 (87%) |
|  | Synonymous | 17363 (8) | 0.99 (0.05) | 15030 (86%) |
|  | Stop-gain/loss | 23 (<1) | 1 (0.01) | 17 (39%) |
|  | Splicing | 5 (<1) | 1 (0) | 5 (100%) |
|  | UTR3/UTR5 | 9943 (5) | 0.99 (0.04) | 8475 (85%) |
|  | ncRNA | 73 (<1) | 0.99 (0.03) | 63 (86%) |

<sup>a</sup> The percentages are in relation to the number of SNPs within the annotation category.
<sup>b</sup> The percentages are in relation to the number of SNPs within the annotation category in RNA.

### Specificity of RNA-seq SNPs

To calculate specificity of our VAP methodology, we focused on variants in coding regions to allow for fair comparison between RNA-seq and WGS data. Approximately 66% of the coding variants identified by WGS were discovered using RNA-seq alone (Fig 6). Given that RNA-seq required less sequencing effort and computational requirements (e.g. 234 million for RNA-seq compared to the 482 million for WGS sequencing reads used in our case study). Using RNA-seq data is advantageous because it enriches for expressed genic regions compared to WGS and therefore will increase the power to detect functionally important SNPs impacting protein sequence.

**Fig 6.**
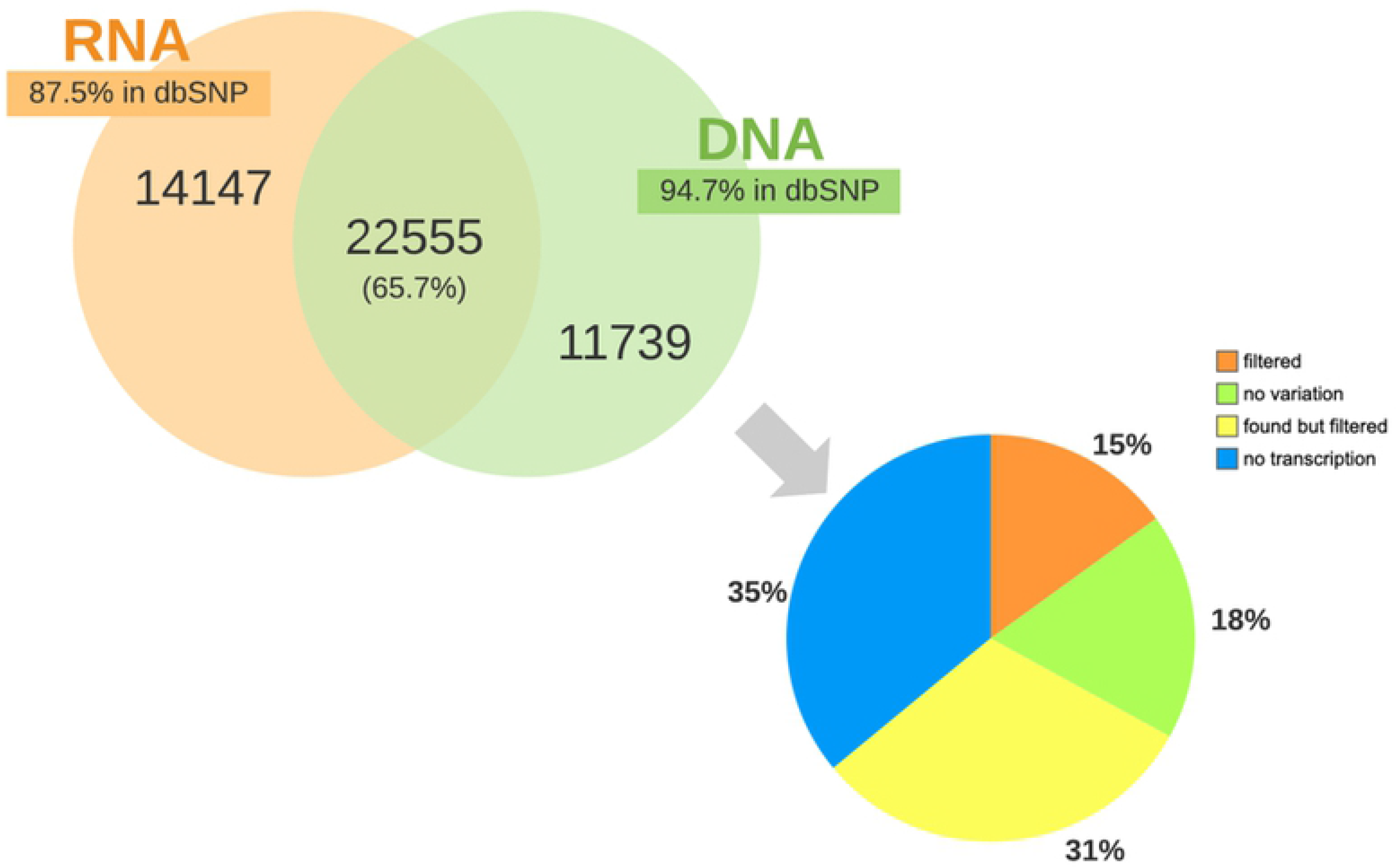
Overlap of SNPs found in coding regions from RNA-seq and WGS. 66% of the coding variants identified in WGS data were found in RNA-seq. However, the remaining WGS coding variants were not detected as a result of either: lack of expression/transcription (“no transcription”), the position was homozygous in RNA (“no variation”), “found but filtered” signifying that the position was detected but removed by one of our filtering steps, or “filtered” which indicates the position was heterozygous but filtered because it didn’t meet the default parameters for variant detection.

We then compared the RNA-seq SNPs in expressed genes (having FPKM > 0.1), and the specificity increased from 66% to over 82% (Fig 7). This shows that a large fraction of genes are expressed at very low levels (Fig 8). Overall the results prove our methodology can achieve high specificity for variant calling in expressed regions of the genome.

**Fig 7.**
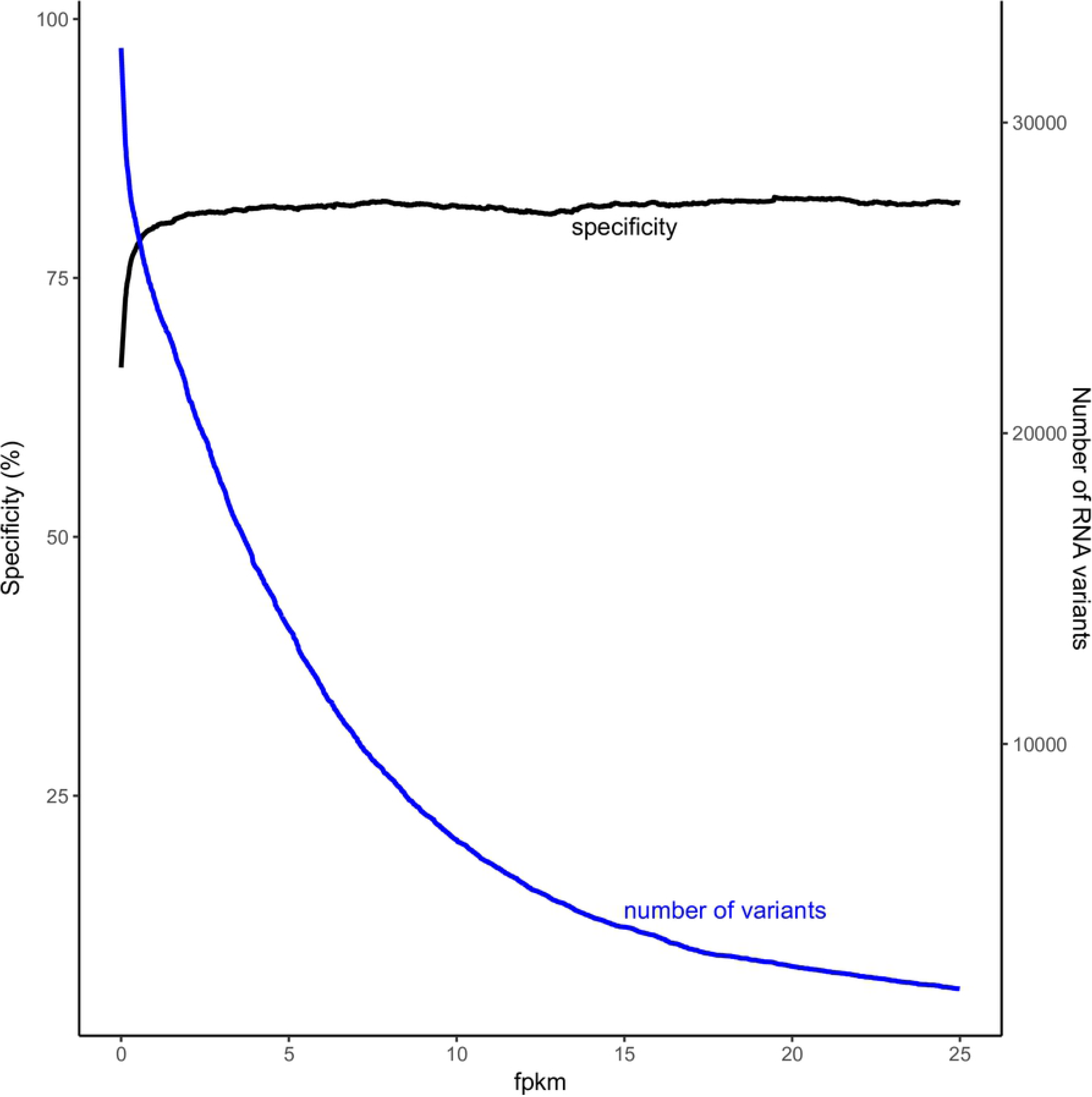
Specificity and number of RNA-seq SNPs detected in relation to the genes expressed (FPKM values).

**Fig 8.**
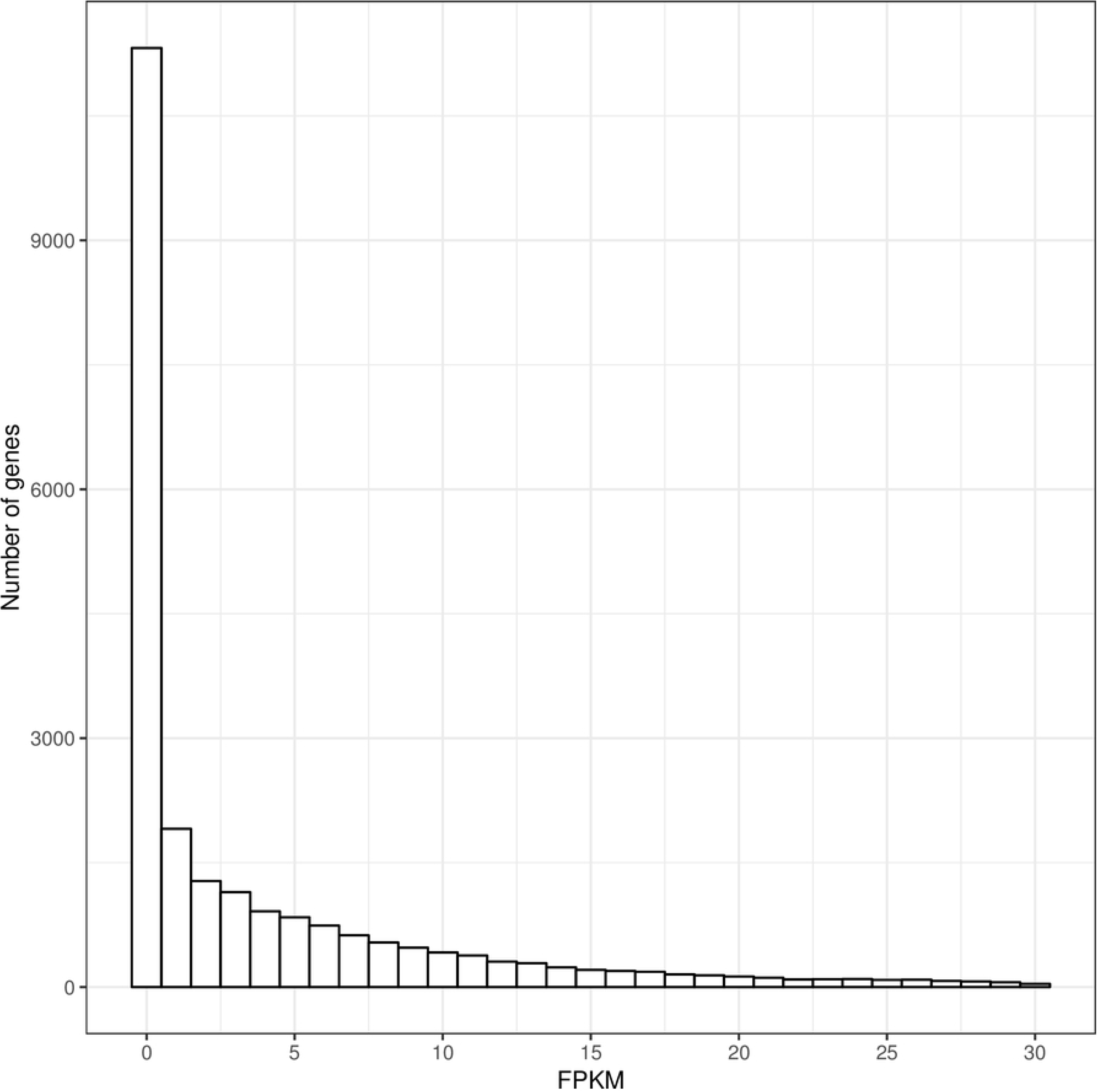
Distribution of expression levels for genes with RNA-seq SNPs.

### Comparison of RNA-seq and 600k Genotyping Panel SNPs

Given the high accuracy of genotyping arrays for SNP discovery, we compared our initially verified RNA-seq SNPs with the genotyped chromosomes identified in the 600k chicken genotyping panel (i.e. the autosomes (GGA1 – 33). A low percentage (10%) of our RNA-seq SNPs overlap with the 600k SNPs (Fig 9), which is largely due to the limitation in the number of variants the genotyping panel is able to capture across different samples. However, 99.9% of the genotyping SNPs were found in dbSNP, proving dbSNP is an adequate method for *in silico* verification of our RNA-seq SNPs.

**Fig 9.**
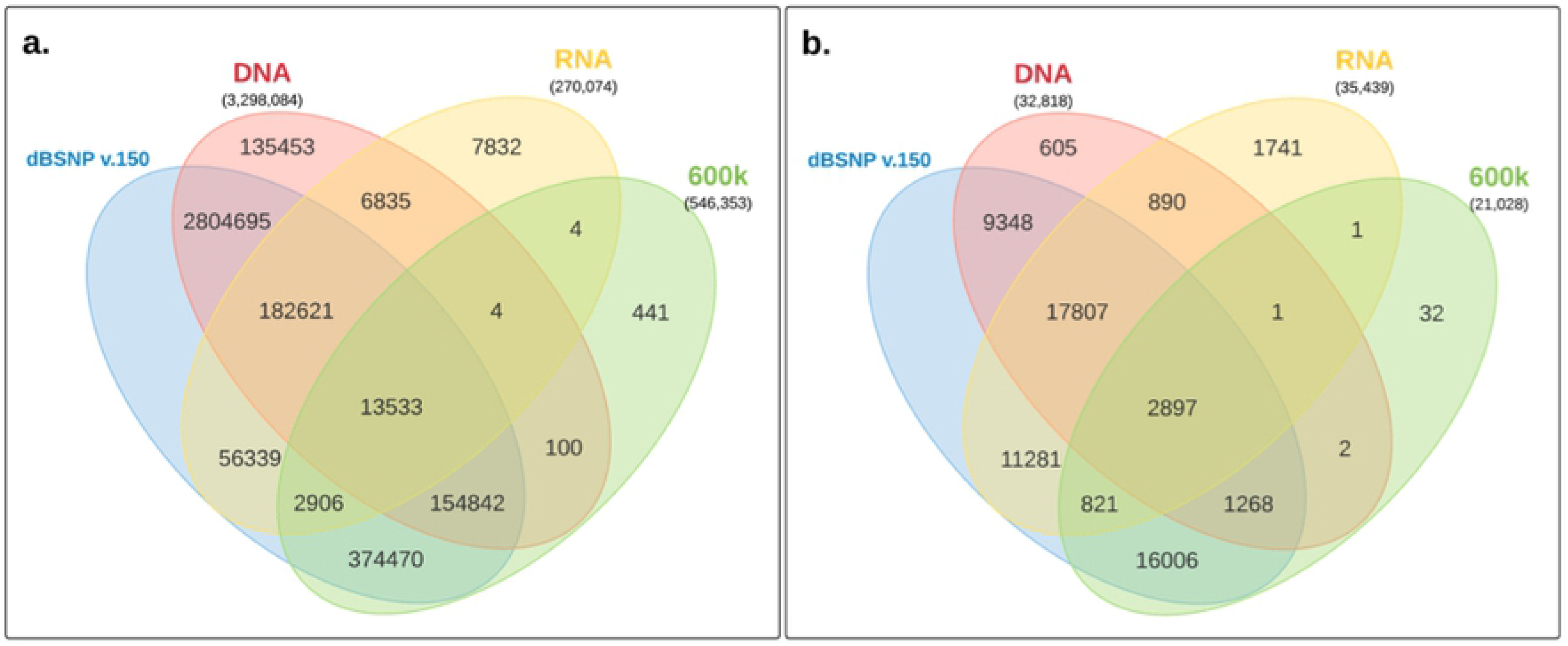
Comparison of SNP calls between 600k Genotyping panel, RNA-seq SNPs, WGS SNPs and dbSNP v150. (a) all autosomal SNPs and (b) autosomal SNPs found in exons.

### RNA–DNA differences (RDD) sites

As mentioned before, our RNA-seq SNPs were notably contributed from transitions which may be attributed to mRNA editing. Further classifications of the RNA-seq SNPs detected in exons reveal 34% of the exonic SNPs verified by dbSNP were not identified in our WGS data. The majority of the RNA SNPs were not found in WGS because of the mapping and filtering parameters as shown in Table 4. Interestingly, 24% of these SNPs were not found because the alternate nucleotide was not present in the DNA sequence potentially indicating RNA–DNA differences (RDD). Consequently, these RDD sites may result from post-transcriptional modification of the RNA sequence, such as RNA editing or alternative splicing.

**Table 4.** Explanation for the 14,147 RNA SNPs not found in WGS data.

| Reason for absence | Number of SNPs (%) |
| --- | --- |
| Position was heterozygous in WGS but filtered because it didn't meet the default parameters for variant detection. | 1225 (8.7) |
| No reads were mapped to region/position. | 1693 (12) |
| Position was homozygous in WGS | 3471 (24.5) |
| Position was heterozygous in WGS but removed by our custom filtering pipeline | 7758 (54.8) |

RNA editing is the most prevalent form of post-transcriptional maturation processes that contributes to transcriptome diversity. It involves the modification of specific nucleotides in the RNA sequence without altering its template DNA (28,32). From our dataset, we identified the three non-synonymous RDD mutations on *CYFIP2*, *GRIA2* and *COG3* previously validated by Frésand et al. in chicken embryos(28) (Table 5). This demonstrates the VAP methodology ability to detect conserved RNA editing phenomena and that it can be used in further discovery of novel post-transcriptional editing events.

**Table 5.** Potentially functional RDD Candidates found in Fayoumi.

| CHROM | POSITION | DNA<br>NUCLEOTIDE | RDD<br>NUCLEOTIDE | AMINO<br>ACID<br>CHANGE | GENE<br>SHORT<br>NAME | VAF<br>SCORE |
| --- | --- | --- | --- | --- | --- | --- |
| chr 1 | 167798513 | A | G | I/V | COG3 | 0.524 |
| chr 4 | 21653669 | A | G | R/G | GRIA2 | 0.703 |
| chr 13 | 11398088 | T | C | K/E | CYFIP2 | 0.375 |

## DISCUSSION

RNA-seq is instrumental in understanding the complexity of the transcriptome. Several methodologies have provided approaches to understanding the varied aspects occurring in the transcriptome, but little has been done in its application to identifying variants in functional regions of the genome. To this aim, we designed the VAP workflow, a multi-aligner strategy using a combination of splice-aware RNA-seq reference mapping tools, variant identification using GATK, and subsequent filtering that allows accurate identification of genomic variants from transcriptome sequencing. Our results show very high precision, sensitivity and specificity, though limited to SNPs occurring in transcribed regions.

Considering the mapping phase of RNA-seq reads is a crucial step in variant calling, we devised a reference mapping strategy using three RNA-seq splice-aware aligners to reduce the prevalence of false positives. The use of the splice-aware aligner allows for accurate assembly of reads because it makes use of both the genome and transcriptome information simultaneously for read mapping.

The ability to call variants from RNA-seq has numerous applications. It enables validation of variants detected by genome sequencing. It also uncovers potential post-transcriptional modifications for gene regulation (Table 5) and allows for detection of previously unidentified variants that may be functionally important but difficult to capture using DNA sequencing or exome sequencing at lower cost. Although our WGS data was not sequenced from the same samples that gave rise to the RNA-seq data, this could explain the poor overlap in our datasets, for instance, 87.5% of RNA-seq variants in exons were not found in WGS though well characterized in dbSNP (Fig 6). Therefore, RNA variants can be used in identifying genetic markers for genetic mapping of traits of interest, thus offering a better understanding of the relationship between genotype and phenotype.

Our VAP methodology shows high precision in calling SNPs from RNA-seq data. It is however limited by the RNA-seq experiments; RNA SNPs are detected only on the transcripts expressed. Regardless of comprehensive coverage, variant detection in some portions of the genome are not guaranteed by RNA-seq because of the potential lack of expression. Also, allele-specific gene expression or tissue-specific gene expression might hamper the discovery of genomic variants given that the allele carrying the variant might not be expressed or the tissues collected might not express the genes of interest.

SNP genotyping offers a highly accurate and alternative method of SNP discovery, and thus offers an additional *in silico* method of validation of our RNA-seq SNPs. However, a low overlap with the 600K chicken genotyping panel was observed (Fig 9). This low overlap is most likely due to the limitations in genotyping panels currently available for any given organism. The genotyping panels are limited by the number of variants they are able to capture across different genetic backgrounds (22). Not surprisingly, the majority of the 600K genotyping variants were also identified in dbSNP, proving that dbSNP an excellent choice for *in silico* validation.

Nevertheless, VAP allows the detection of variants even for lowly expressed genes. To obtain higher confidence in variant calls, pooling multiple data sets (i.e. RNA-seq from different tissues) can increase the coverage thereby facilitate variant discovery in regions of interest that would have otherwise been missed. Our study demonstrates that variants calling from RNA-seq experiments can tremendously benefit from an increased number of reads increasing the coverage of genomic regions especially for whole genome analysis; nevertheless even our small sample size allowed for reliable calling of variants and enriching for variants in exonic regions.

Despite the limitations of calling genomic variants from RNA-seq data, our work shows high sensitivity and specificity in SNP calls from RNA-seq data. SNP calling from RNA-seq will not replace WGS or exome-sequencing (WES) approaches but rather offers a suitable alternative to either approaches and might complement or be used to validate SNPs detected from either WGS or WES. Overall, we present a valuable methodology that provides an avenue to analyze genomic SNPs from RNA-seq data alone.

## AUTHOR CONTRIBUTIONS

M.O.A. designed the methodology, performed the data analysis and drafted the manuscript. C.J.S. secured the funding, advised in the research and revised the manuscript. S.J.L. and B.A. provided the data and revised the manuscript.

